# Evidence for the digital programming of cellular differentiation: characterization of binary switches in lineage-determining transcription factors

**DOI:** 10.1101/2022.06.22.497234

**Authors:** Hongchuan Li, Md Ahasanur Rahman, Paul W Wright, Jennie Cao, Shashikala Ratnayake, Qingrong Chen, Chunhua Yan, Daoud Meerzaman, Aharon G Freud, Stephen K Anderson

**Author notes:** The content of this publication does not necessarily reflect the views or policies of the Department of Health and Human Services, nor does mention of trade names, commercial products, or organizations imply endorsement by the US Government.

## Abstract

Previous studies of the murine *Ly49* and human *KIR* gene clusters have revealed a role for bidirectional promoters in the control of variegated gene expression. Whether or not competing promoters control other instances of cell fate determination remains an outstanding question. Although divergent transcripts within 300 bp are found in ~6% of human genes, an analysis of human transcription factor (TF) genes (1640) revealed that 33% possess stable divergent transcripts with a transcription start site (TSS) less than 300 bp upstream, with many separated by less than 30 bp, indicating an enrichment for potential binary switches in TFs. We have performed a detailed examination of putative bidirectional promoter switches in three lineage-determining TF genes: *AHR, GATA3*, and *RORγT*. These genes also contain additional pairs of opposing promoters that would prevent simultaneous transcription of sense and antisense, and thus may represent simple on/off switches rather than probabilistic switches. In situ RNA hybridization of human tissues revealed mutually exclusive expression of sense versus antisense transcription, indicating switching between stable sense or antisense transcriptional states. Single-cell RNAseq confirmed the separate sense/antisense states, and revealed the identity of cells with active switch elements. Differential gene expression analysis of cells with antisense switch transcripts revealed an enrichment for genes found in immature/stem cells and a lack of genes associated with terminally differentiated cells. Taken together, these data indicate that there is a digital component to the differentiation program mediated by binary promoter switches in lineage-determining transcription factors.

## Introduction

Soon after the completion of the human genome project, several aspects of the human genome became apparent: the small percentage of the genome that codes for protein (~2%), the abundance of non-coding RNA transcripts (70% of the genome is transcribed), and the significant frequency of divergent transcripts controlled by a central bidirectional promoter region (ENCODE Project Consortium; Trinklein et al., 2004). Approximately 12% of the transcription start sites of human genes are located on opposite strands within 1 kb of each other, and were thus defined as bidirectional promoters with putative shared regulatory elements controlling the divergent transcription of linked genes. Furthermore, 6% of human genes are arranged in divergent pairs separated by less than 300 bp, raising the possibility of competing regulatory elements, or perhaps binary switches. The majority of bidirectional promoters may mediate coordinated regulation of the linked genes, since sense and antisense transcription was highly correlated in 89% of divergent transcript pairs. A negative correlation between the expression of divergent transcripts was observed in 11% of the bidirectional promoters analyzed, indicating potential switch-like behavior (Trinklein et al., 2004).

More recent studies have revealed that transcription of both DNA strands occurs in most enhancers and promoters. GRO-seq of exonuclease-deficient cells has enabled the detection of short, rapidly degraded transcripts from the antisense strand of promoters previously described as unidirectional (Core et al., 2008). This may reflect the opportunistic binding and entry of a second RNA polymerase complex to the open, nucleosome-depleted region (NDR) that is created upstream from a RNA polII complex that has initiated transcription, but is stalled. The lack of splice donor sites and the presence of polyA-addition signals is associated with the rapid degradation of enhancer RNAs and antisense transcripts from gene promoters (Andersson and Sandelin, 2020). However, efficient bidirectional transcription of mRNA from the paired arrangement of genes is associated with higher rates of transcription, indicating that the head to head arrangement of housekeeping genes on either side of a GC-rich promoter region allows for maximal production of transcripts from essential genes (Adachi and Lieber, 2002).

Closely-spaced divergent promoter pairs may act as binary switches and play a role in the programming of cell fate or the generation of diverse phenotypes within a given cell type. The subset of promoters capable of acting as switches is likely quite small, since a negative correlation between the expression of sense and antisense divergent transcripts was identified in approximately 1% of human genes. The ability of a bidirectional promoter to function as a probabilistic switch was demonstrated in the murine *Ly49* gene cluster, a family of MHC class I receptors that are stochastically expressed by NK cells (Saleh et al., 2004). A 119 bp core bidirectional promoter was able to generate divergent transcripts, but for an individual *Ly49* gene, only one direction was active at any given time, allowing it to act as a binary switch. Competition of transcription factors for overlapping binding sites determined the relative strength of the competing promoters and the probability of transcription in a given direction. The relative strength of the sense versus antisense promoters in each inhibitory *Ly49* gene correlated with the percentage of natural killer cells expressing that receptor, supporting a promoter competition model of selective gene activation.

We have conducted in silico analyses of lineage-determining transcription factors, and discovered that many of them possess spliced polyadenylated antisense transcripts initiated in close proximity to the transcription start site of the gene. This suggests that they may function as probabilistic binary switches and generate a variegated expression pattern of these transcription factors. A defining characteristic of a binary promoter switch is the presence of mutually exclusive transcriptional states where only the sense or antisense transcript is generated over a sustained period of time. We have examined the potential for switch behavior in the promoters of three lineage-determining transcription factors: RORγT; GATA3; AHR. Our previous studies demonstrated switch behavior using fluorescent reporter constructs, however recent technological advances have provided additional methods to assay bidirectional transcription from genes in their native context. 5’-directed single-cell RNAseq allows the quantitation of sense and antisense in a single cell, and can identify transcription factors associated with activity in a given direction. Improvements in the sensitivity of in situ RNA hybridization allows direct detection of individual sense and antisense transcripts in intact tissue. Single-cell RNAseq and in situ RNA hybridization studies of sense and antisense transcripts from the human *RORγT, GATA3*, and *AHR* gene promoters reveals that sense and antisense transcripts are mutually exclusive, indicating that the expression of these transcription factors is controlled by binary promoter switches.

## Results

### The RORγT promoter is bidirectional and produces spliced antisense transcripts

RORC, or RAR-related Orphan Receptor C, belongs to a family of hormone nuclear receptors that bind to specific ROR-response elements, which consist of a core AGGTCA motif preceded by a 5-bp A/T rich sequence (Jetten et al., 2001; Capone and Volpe, 2020). The *RORC* gene produces two protein isoforms, RORγ and RORγT, transcribed from alternative promoters. RORγ expression has been detected in many tissues, including liver, kidney, lung, muscle, and heart (He et al., 1998; Ruan et al., 2011; Eberl et al., 2003). RORγT, thus named due to its high expression in the thymus, is the master regulator of Th17 differentiation (Ivanov et al., 2006). Examination of the *RORγT* promoter region in the USC Genome Browser (http://genome.ucsc.edu) revealed the presence of a spliced antisense transcript initiating 192 bp upstream from the *RORγT* transcription start site. An additional antisense transcript originating in the first *RORC* intron, opposing the *RORC* promoter was also observed (Figure 1A).

**Figure 1.**
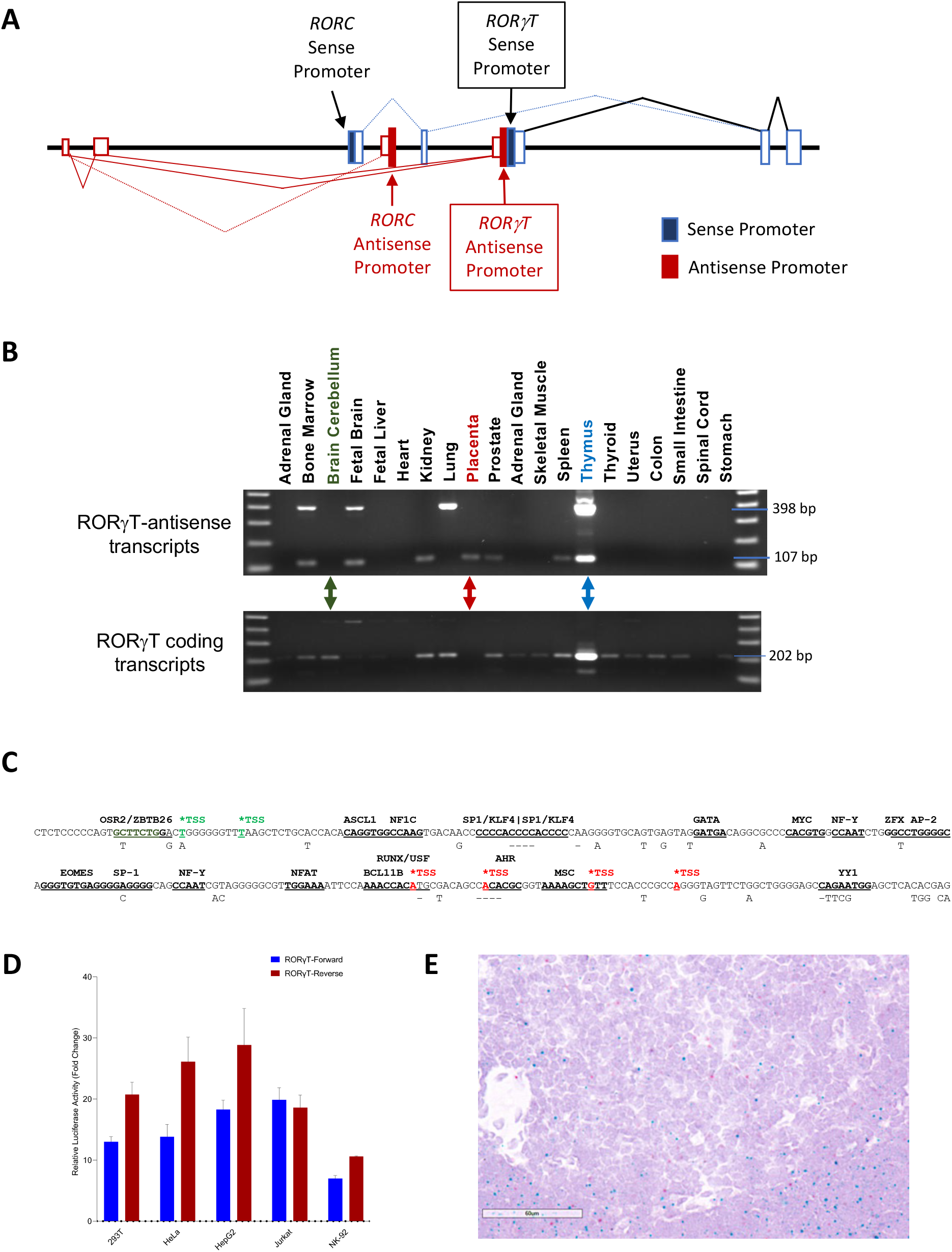
The *RORC* gene contains a potential switch region. (A) Schematic representation of the 5’ region of the *RORC* gene is shown. Open blue rectangles represent coding strand exons, and red open rectangles show antisense exons. Exon splicing patterns are indicated by lines joining the exons. Filled boxes indicate promoter regions. Labeled boxes with arrows indicate the location of the adjacent *RORγT* promoters producing divergent transcripts. (B) A cDNA panel of 20 human tissues was subject to PCR with primers specific for either RORγT antisense (upper panel) or coding strand transcripts. Examples of tissues that contain only sense (green type), only antisense (red type), or both (blue type) are indicated by the colored arrows. (C) The sequence of the human *RORC* gene in the region spanning divergent sense and antisense TSS is shown. The coding strand TSS is indicated by the green asterisk, and the multiple antisense TSS are marked by red asterisks. Predicted TF-binding sites are indicated by bold underlined type. Only differing nucleotides present in the mouse *RORC* gene are shown below the human sequence. (D) The relative luciferase activity of a 278 bp *RORγT* bidirectional promoter fragment in five cell lines. Fold change in light units relative to empty pGL3 vector is shown on the y-axis. Values represent the mean and error bars indicate the SEM of at least three independent experiments. Promoter activity from either *RORγT-* Forward (coding strand; blue) or *RORγT*-Reverse (antisense; red) orientations of the promoter fragment are shown for each cell line. (E) In situ RNA hybridization with RNAscope probes was performed on human thymus tissue. The red spots indicate binding of a probe that recognizes a 138 bp region in the first exon of the RORγT antisense transcript. The blue spots contain a probe that binds to a 104 bp region contained within the first exon of the RORγT mRNA.

In order to determine if the sense and antisense transcripts were coordinately regulated, RT-PCR with primers specific for either the sense or antisense *RORγT* transcripts was performed on a panel of human tissue RNAs, demonstrating the presence of sense and antisense transcripts in multiple tissues, with the highest expression of both transcripts observed in thymus (Figure 1B). Interestingly, the ratio of sense to antisense varied among tissues, and several tissues exhibited only sense activity, while placenta only had antisense transcripts, indicating that sense and antisense activity were not inextricably linked, supporting the possibility of promoter switching.

Examination of the *RORγT* bidirectional promoter region revealed an abundance of predicted transcription factor binding sites, including sites for other lineage-determining transcription factors, AHR, GATA, and EOMES (Figure 1C). In addition, a group of transcription factor-binding motifs that are enriched in bidirectional promoters: MYC, NF-Y, and YY1 are also present (Lin et al., 2007). This promoter region is highly conserved among vertebrates, and comparison of the human and mouse genes reveals 89% nucleotide identity, with most of the predicted TF-binding sites being conserved (Figure 1C).

A 278 bp fragment containing the bidirectional promoter region was cloned into the pGL3 luciferase reporter vector in both the sense and antisense orientations. Figure 1D shows the relative promoter activity of the *RORγT* bidirectional promoter in several cell lines. Antisense activity was dominant in the human embryonic kidney line HEK293T, human cervical cancer cell line HeLa, human liver HepG2 cell line, and human NK92 NK cell line, while sense/antisense activity was similar in the human T cell Jurkat cell line. These observations suggest that differences in transcription factor expression in individual cell lines affects competition between transcription factors driving either sense or antisense transcription.

In order to determine if the *RORγT* bidirectional promoter could function as a stable binary switch, in situ RNA hybridization was performed on human thymus tissue which expresses high levels of both sense and antisense transcripts (Figure 1B). The expression of sense and antisense *RORγT* transcripts were primarily independently expressed, with only rare instances in which both were detected in a single cell (Figure 1E).

### The AHR gene contains a bidirectional promoter and adjacent opposing promoters

The aryl hydrocarbon receptor (AHR), a member of the basic helix-loop-helix-PAS ligand-activated transcription factor family, is responsible for regulating hypoxia, circadian rhythms, and other cellular processes such as differentiation and apoptosis, through binding to the aryl hydrocarbon response element (AHRE), 5’-GCGTG-3’ (Garrison et al., 2000; Vogel et al., 2014; Scoville et al., 2018; Racky et al., 2004). Originally discovered for its role in sensing environmental toxins such as 2,3,7,8-tetrachlorodibenzo-p-dioxin (TCDD), various additional exogenous ligands have been identified. However, kynurenine, the first breakdown product in the IDO-dependent tryptophan degradation pathway, has been identified as an endogenous ligand that activates AHR (Harper et al., 2006; Mezrich et al., 2010).

Visualization of the *AHR* gene in the USC Genome Browser and the Eukaryotic Promoter Database (EPD; Dreos et al. 2015) revealed two coding strand promoters initiating transcription either 27bp (EPD: AHR_7) or 613 bp (EPD: AHR_1) upstream from the AHR translation initiation site (Figure 2A). There are multiple antisense transcription start sites (TSS) contained within the first *AHR* exon and 5’ region of the intron, with two major sites located 53 bp downstream from the AHR start codon and 34 bp downstream from the intron-exon junction. This indicates that an antisense promoter (asPro1; Figure 2A) is present in the first intron that opposes transcription from the *AHR* coding strand promoters. An additional antisense TSS is located 205 bp upstream from the AHR_1 transcript initiation site, indicating bidirectional activity in this region indicated by asPro2 in Figure 2A. Therefore, the *AHR* gene contains both overlapping and closely-spaced divergent transcripts associated with two coding-strand promoters. Both antisense transcripts generate spliced, polyadenylated transcripts lacking significant open reading frames (Figure 2A). RT-PCR of a panel of human tissue RNAs with primers specific for asPro1 or asPro2 antisense *AHR* transcripts demonstrated the presence of antisense transcripts in multiple tissues, with the highest expression of the Pro2 antisense transcript observed in heart (Figure 2B). The AHR-coding transcript was detected in all tissues, however, it was not possible to distinguish between transcripts originating from either the AHR_1 or AHR_7 promoters due to the overlapping nature of these transcripts. Multiple tissues lacked Pro2 antisense expression as observed for the *RORγT* bidirectional element, indicating potential switch behavior. In contrast, the Pro1 antisense transcript was detected in all tissues with differing intensities, suggesting differential activity or variation in the percentage of cells expressing this antisense in each tissue.

**Figure 2.**
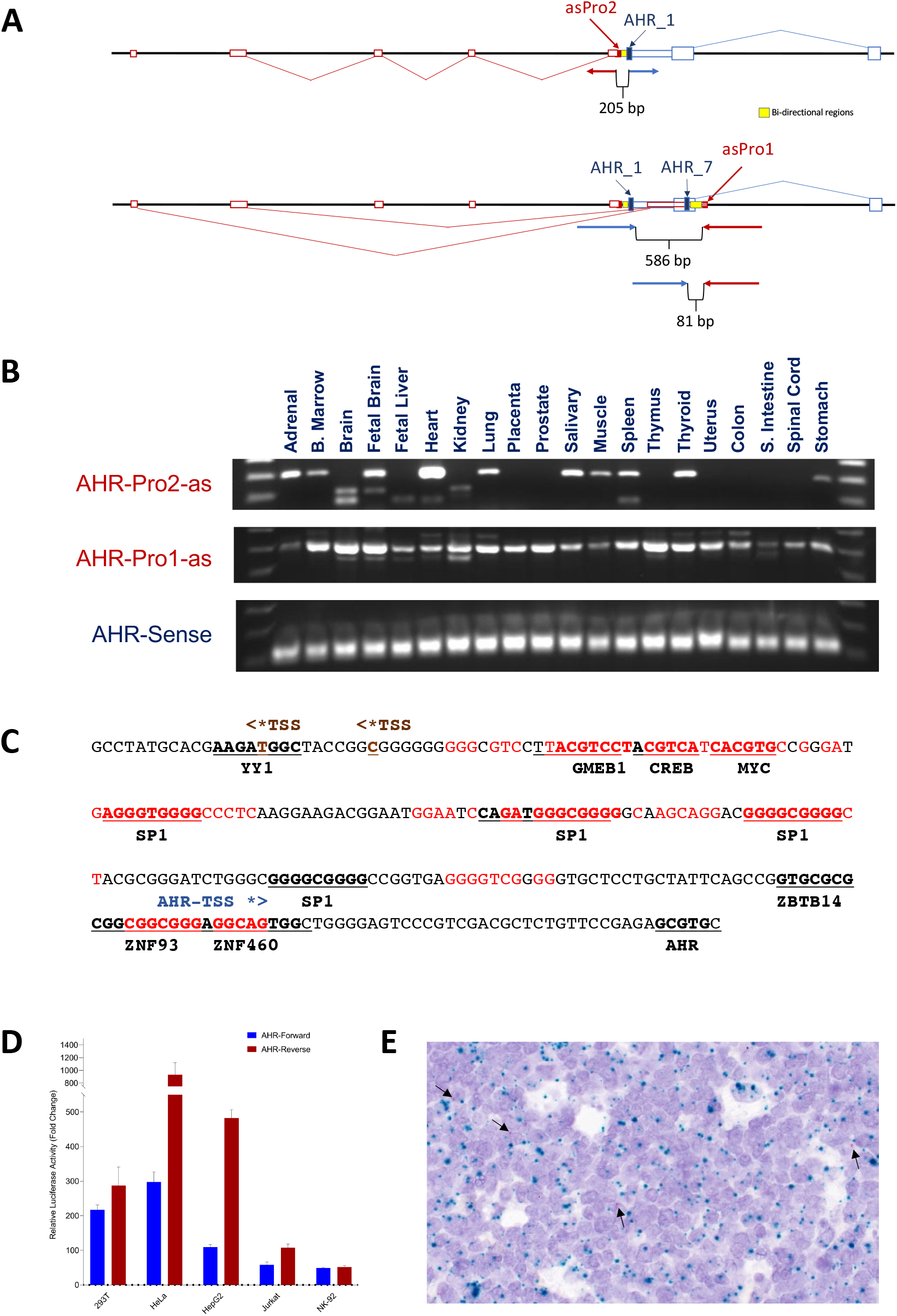
The *AHR* gene contains divergent and opposing bidirectional switch regions. (A) Schematic representation of the 5’ region of the *AHR* gene is shown. Open blue boxes represent coding strand exons, with larger blue boxes indicating the open reading frame, and red open boxes show antisense exons. Exon splicing patterns are indicated by lines joining the exons. Filled boxes indicate promoter regions. The upper panel shows the divergent transcripts produced by asPro2 and the AHR_1 promoter. The labeled bracket below the bidirectional promoter region indicates the separation between divergent TSS. The lower panel shows the opposing transcripts generated by the AHR_1/AHR_7 sense strand promoters and the antisense transcript from asPro1. The labeled brackets beneath the promoter regions indicate the separation between either AHR_1 and asPro1 TSS (586 bp), or AHR_7 and asPro1 TSS (81 bp). (*B*) A cDNA panel of 20 human tissues was subject to PCR with primers specific for AHR Pro2 antisense (AHR-Pro2-as; upper panel), AHR Pro1 antisense (AHR-Pro1-as; middle panel) or coding strand transcripts originating from either the AHR_1 or AHR_7 promoters (AHR-Sense; lower panel). (*C*) The sequence of the human *AHR* gene in the region spanning divergent AHR_1-sense and antisense Pro2 TSS is shown. The coding strand TSS is indicated by the blue asterisk, and two antisense TSS are marked by brown asterisks. Predicted TF-binding sites are indicated by bold underlined type. Nucleotides that are conserved in the mouse *AHR* gene are shown in red type. (*D*) The relative luciferase activity of a 264 bp *AHR* bidirectional promoter fragment in five cell lines is shown. Fold change in light units relative to empty pGL3 vector is shown on the y-axis. Values represent the mean and error bars indicate the SEM of at least three independent experiments. Promoter activity from either forward (AHR_1; blue) or reverse (asPro2; red) orientations of the bidirectional promoter fragment are shown for each cell line. (*E*). In situ RNA hybridization with AHR-Pro1 and asPro2-specific probes was performed on human tonsil tissue. The red spots, indicated by the black arrows, represent binding of a probe that recognizes a 39 bp region in the first exon of the asPro2 antisense transcript. The blue spots correspond to a probe that binds to a 276 bp region contained within the first exon of the Pro1-derived AHR mRNA.

The 264 bp sequence of the asPro2/AHR_1 bidirectional promoter region shown in Figure 2C also contains predicted SP1, MYC, and YY1 sites, as observed for the *RORγT* switch element. Of particular interest, this element contains a predicted dioxin-response factor, ZBTB14, binding site 15 bp upstream of the sense strand TSS, and an AHR-binding site 37 bp downstream, suggesting possible feedback modulation of the coding strand promoter. In addition, there is a binding site for ZNF93, a transcription factor associated with the silencing of retrotransposons in stem cells (Jacobs et al., 2014). In contrast to the sequence conservation of the entire *RORγT* element in the mouse, *AHR* has several regions of sequence homology, with conservation of the ZNF93 and SP1-binding sites, and a homologous block of tandem GMEB1/CREB/MYC sites. A 264 bp fragment of the asPro2/AHR_1 promoter region was cloned into pGL3 in both orientations, and the relative forward/reverse activity is shown in Figure 2D. Antisense promoter activity was dominant in four of the five cell lines, in contrast to the high expression of sense transcripts observed in most of the human tissues tested (Figure 2C), indicating differential activity in immortalized cell lines or that additional sequence elements in the *AHR* gene or the downstream AHR_7 promoter may be required for optimal forward promoter activity.

In situ RNA hybridization with a probe specific for the asPro2 transcript was performed on human heart tissue, which had the highest level of antisense expression of the tissues examined (Figure 2C). Although sense transcript was abundantly expressed, consistent with the RT-PCR results obtained with human tissues, antisense was only detected in a few cells. Additional human tissues were tested, including thyroid, lymph node, and tonsil. Weak expression of the asPro2 transcript was detected in a small number of cells in tonsil tissue, indicating that antisense transcription may be a rare event associated with differentiating cells, or the antisense transcript is highly unstable (Figure 2E).

The presence of an opposing promoter in the first intron of the *AHR* gene (asPro1), potentially creating an on/off-switch, is intriguing, especially with regard to a recent study that demonstrated the ability of AHR antagonists to enhance ESC differentiation (Chen et al., 2022). If the function of the asPro1 promoter is to downregulate AHR expression in differentiating cells, then its expression should be limited to early progenitors. Single-cell RNAseq has the potential to characterize cells expressing *AHR* asPro1 transcripts. In light of recent studies demonstrating that AHR inhibition favors NK cell differentiation (Lordo et al., 2021), we examined *AHR* transcript expression in ex vivo human NK cells by single-cell RNAseq. Chromium-5’ V2 mRNA libraries were generated using purified NK cells obtained from four donors. More than 30,000 cells were sequenced, and individual sequence reads were analyzed using the Integrated Genome Viewer (IGV; Robinson et al., 2011). Analysis of the 5’-region of the *AHR* gene revealed a lack of spliced asPro2 transcripts, however a significant number of asPro1 transcripts were observed (Figure 3A).

**Figure 3.**
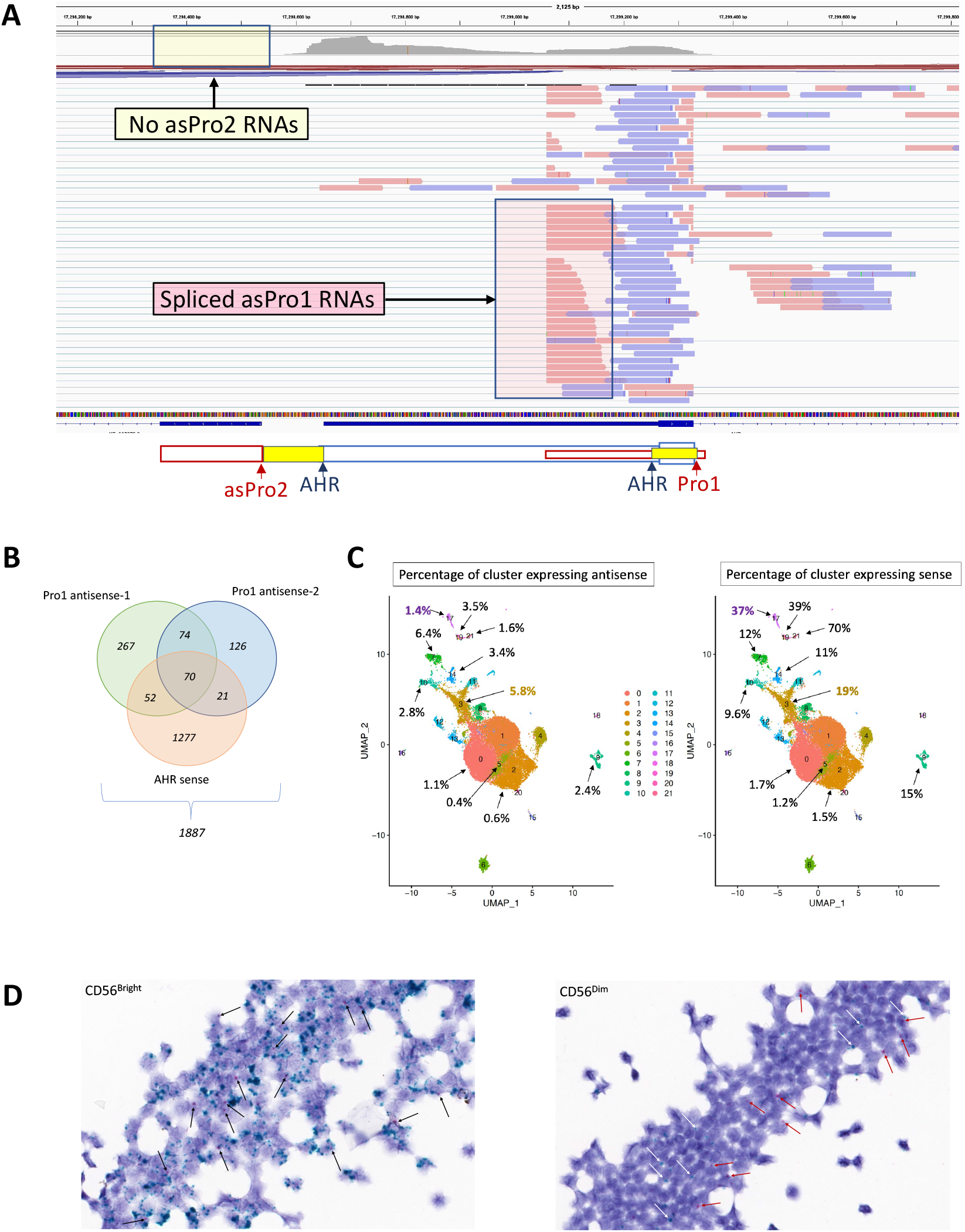
AHR asPro1 activity varies with cell type and NK cell differentiation. (*A*) An image taken from an IGV analysis of RNAseq reads in the AHR 5’-region is shown. A schematic of the location of the sense and antisense promoters is shown below the image. The region highlighted by the pink box represents spliced RNAs transcribed from the asPro1 promoter. The empty yellow box in the sequence coverage track at the top of the image shows the region where asPro2 RNAs would be detected if present. (*B*) A Venn diagram summarizing the expression of AHR sense and antisense transcripts in individual cells is shown. The two major antisense splice forms (Pro1 antisense-1 and Pro1 antisense-2) are shown together with AHR sense transcripts. (*C*). A UMAP generated with 35PCs at a resolution of 0.3 utilizing 157864 features across 30637 samples is shown. The percentage of cells within a cluster expressing AHR antisense (left panel) or AHR sense (right panel) is indicated for selected cluster. (*D*) In situ RNA hybridization with AHR-sense and asPro1-specific probes was performed on sorted human CD56^Bright^ (left panel) and CD56^Dim^ (right panel) NK cells. The red spots, indicated by the black arrows, represent binding of a probe that recognizes a 149 bp region in the second exon of the asPro1 antisense transcript. The blue spots correspond to a probe that binds to a 276 bp region contained within the first exon of the Pro1-derived AHR mRNA.

Bioinformatic analysis of single cells expressing *AHR* sense or Pro1 antisense transcripts indicated that 6% of the total cell population expressed *AHR* transcripts (1887/30637; Figure 3B). Of the AHR-expressing cells, 68% expressed sense transcripts only, 25% expressed antisense only, and 7.5% expressed both. This is significantly less than the predicted 17% (.68 x .25) co-expression if the sense and antisense are independently expressed. However, if both *AHR* alleles are active in a given cell, then the percentage per allele would be 34% and 12.5%, resulting in a predicted co-expression of 4%, closer to the observed percentage of coexpression. These observations are consistent with mutually exclusive expression of the opposing sense and antisense transcripts, with rare instances of co-expression occurring due to antisense and sense being produced simultaneously from different alleles in the same cell. To further explore the phenotype of the cells expressing *AHR* transcripts, a UMAP of the single cells was generated and the expression of sense or antisense *AHR* in distinct clusters was analyzed (Figure 3C; Supplemental Tables 1 and 2). A rapid purification method was used to limit the time between blood collection and the single cell RNA analysis, in order to obtain data that better reflects the in vivo expression patterns. This resulted in a much larger number of clusters due to the presence of non-NK populations such as monocytes (cluster 9), neutrophil/mast cells (cluster 15) and B cells (cluster 18), but it also captured progenitor cell populations (clusters 17, 19, 21) and multiple NK populations not previously reported. *AHR* sense was highly expressed by progenitor cell populations (clusters 17, 19, and 21), and antisense was preferentially expressed by immature CD56^bright^ NK cells (clusters 3, 7, 14). The CD56^dim^ (clusters 0, 1), activated NK (cluster 5), and terminally differentiated NK cells (cluster 2) had the lowest levels of *AHR* transcripts. Interestingly, the ratio of sense to antisense was very high (>10) in progenitor cells, and decreased in the immature NK populations (2-3 fold sense:antisense). The percentage of *AHR* sense expressing cells also decreased dramatically, from approximately 40% in progenitors, to ~10% in immature NK cells, and less than 2% in mature NK cells. This indicates that AHR is downregulated during differentiation, in line with observations that AHR antagonism increases NK cell differentiation (Lordo et al., 2021; Chen et al., 2022).

In situ RNA hybridization of purified blood CD56^bright^ and CD56^dim^ NK cells was performed to confirm the results obtained by single-cell RNAseq. As shown in Figure 3D, *AHR* sense transcripts were abundantly expressed by CD56^bright^ cells with multiple spots per cell, whereas only a few cells displayed single sense transcripts in the CD56^dim^ population. Antisense transcripts were less abundant, with only one spot per cell in either the bright or dim population. Sense and antisense transcripts were expressed independently in the CD56^dim^ cells, however very few cells in the CD56^bright^ cells were observed to express antisense alone, which could be due the high level of sense transcripts in overlapping/adjacent cells appearing to colocalize with antisense, or alternatively, it could be indicative of a recent switch in *AHR* transcription direction.

In order to determine if a particular cellular phenotype or TF expression pattern was associated with antisense transcription, a differential gene expression analysis was conducted, comparing *AHR* antisense expressing cells with all others. This revealed a group of 74 genes that were significantly upregulated (Table 1; Supplemental Table 3).

**Table 1.**
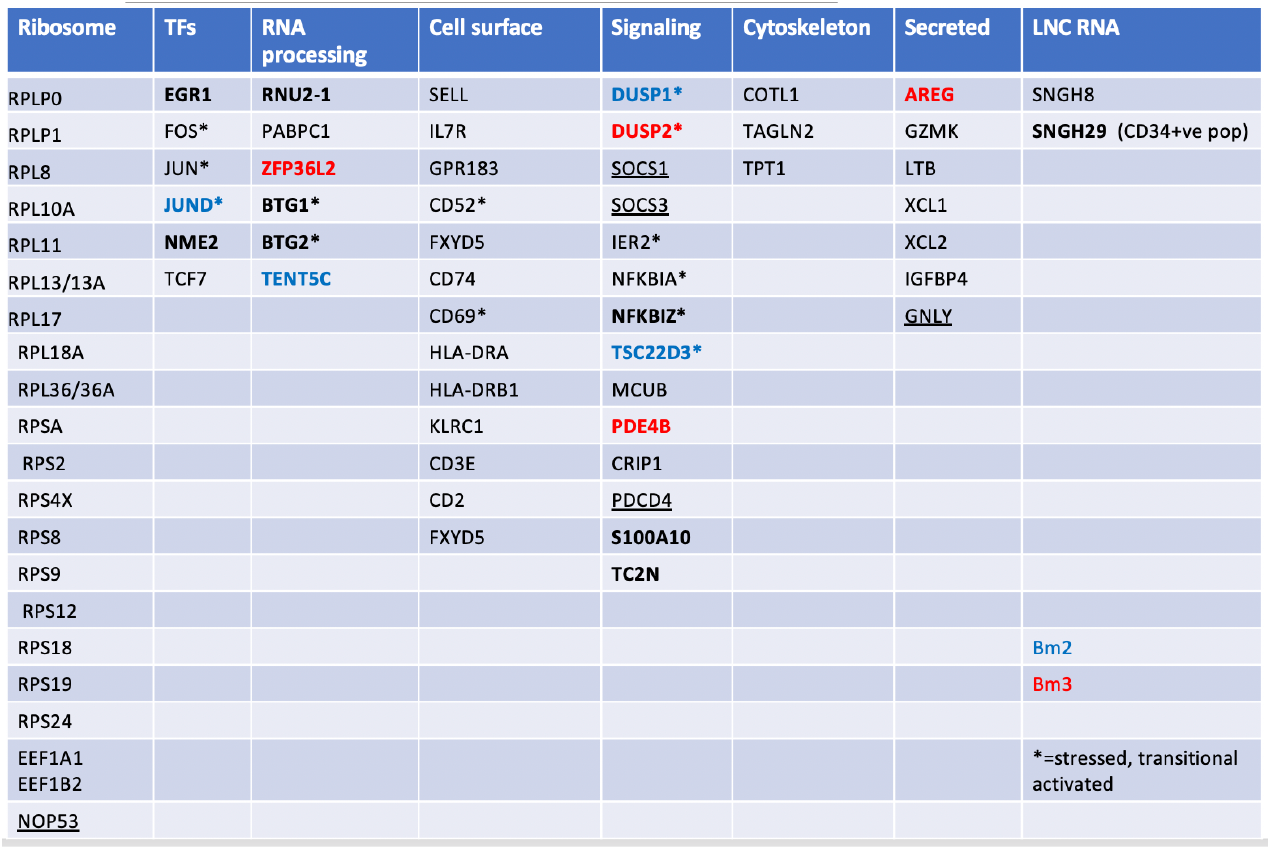
Genes upregulated in cells expressing Pro1 antisense transcripts. The 74 genes with increased expression in cells expressing the AHR antisense are listed in eight functional categories. Most of the listed genes are highly expressed in blood CD56^bright^ NK cells, with exceptions shown in bold type. Underlined genes were not found in previously published blood CD56^bright^ gene lists, but were detected in the CD56^bright^ cluster in the current study. Genes overexpressed in bone marrow CD56^bright^ cells (blue type) or their progenitors (red type) are indicated. Asterisks indicate genes associated with activated, transitional, or stressed NK cells.

The majority of the genes associated with asPro1 transcription correspond to those observed in the CD56^bright^ subset of immature NK cells by previous studies (Crinier et al., 2021; Smith et al., 2020; Yang et al., 2019). Of the 21 genes not previously associated with the blood CD56^bright^ phenotype, 4 were associated with the CD56^bright^ cluster in the current dataset, and 8 were found to be upregulated in CD56^bright^ NK cells and their progenitors from the bone marrow (Crinier et al., 2021), suggesting an association of *AHR* antisense transcription with the immature cell state. The remaining upregulated genes not associated with the CD56^bright^ phenotype belonged to two major classes: activation-associated genes and genes associated with progenitors. The up-regulation of EGR1, FOS, JUN, DUSP1/2, NFKBIA/Z, and SOCS1/3 suggests that the *AHR* antisense is expressed by cells that have received growth factor/cytokine stimulation. Of particular interest is the group of genes associated with mRNA deadenylation (Amine et al., 2021): PABPC1, BTG1, BTG2, and ZFP36L2. This group of genes has been shown to be important for the maintenance of progenitor cells in several systems (Zhang et al., 2013; Tijchon et al., 2016; Hwang et al, 2020). In addition, the transcription factor NME2 has been shown to maintain the stemness of gastric cancer stem-like cells (Qi et al., 2021). An additional differential gene expression analysis was performed in order to determine if there were any genes specifically upregulated in antisense expressing cells as compared to cells containing AHR sense transcripts (Supplemental Table 4). The number of differentially expressed genes associated with *AHR* sense expression was significantly greater (253 genes upregulated) due to the high expression of AHR in several non-NK cell clusters, such as monocytes. Of the genes associated with *AHR* transcription, ZFP36L2, BTG2, TENT5C, and GNLY were the only genes specifically upregulated in antisense expressing cells. The association of genes involved in removal of pA tails (ZFP36L2, BTG2) with the TENT5C gene which encodes a non-canonical polyA RNA polymerase is intriguing, and may indicate that the *AHR* antisense promoter is activated in progenitor cells that are transitioning from a progenitor state to a more mature cell type.

In order to determine if any of the six TFs upregulated in cells expressing *AHR* sense/antisense transcripts could have a direct effect on expression, the sequence of the asPro1 and AHR_7 promoter region was analyzed. Figure 4 shows the sequence of the region containing the *AHR* sense promoter and the asPro1 promoter. No binding sites for FOS/JUN, or TCF7 were present, however there are two predicted EGR1 binding sites 36 bp and 56 bp upstream from the major antisense TSS. This suggests that antisense *AHR* transcription may be triggered by growth factor stimulation through EGR1 activation, which is itself upregulated through FOS/JUN binding sites in the *EGR1* promoter (Wang et al., 2021).

**Figure 4.**
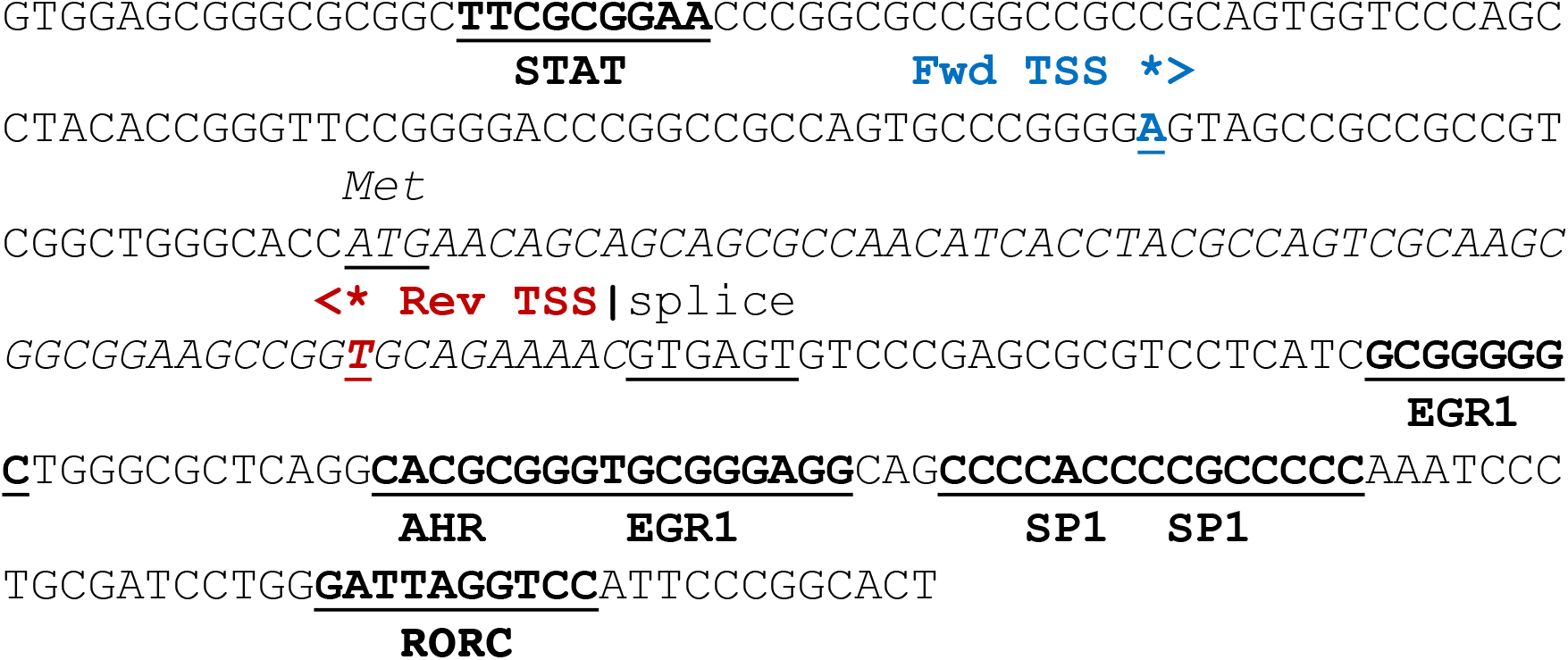
Sequence of the opposing AHR_7/asPro1 promoter region. The sequence of the human *AHR* gene spanning the AHR_7 TSS and the asPro1 TSS is shown. The sense-strand TSS is indicated by the blue asterisk, and the antisense TSS is indicated by a red asterisk. Predicted transcription factor-binding sites are underlined in bold. The AHR coding region is italicized, and the initiation codon and splice donor site are underlined.

### The GATA3 gene contains multiple sense and antisense promoters

GATA3 is a member of a family of zinc-finger transcription factors that bind to the consensus sequence 5’-AGATAA(A/G)-3’. GATA3 is abundantly expressed in various tissues throughout development, including the brain, placenta, and kidney (Grégoire and Roméo, 1999), however, its role in T-cell differentiation has been extensively studied. As the master regulator of Th2 (T helper type 2) lineage commitment, GATA3 suppresses the Th1 pathway while also upregulating the transcription of cytokines IL-4, IL-5, and IL-13 (Tindemans et al., 2014; Scheinman and Avni, 2009).

Examination of the *GATA3* gene in the USC Genome Browser and the EPD revealed the presence of 4 distinct sense transcripts initiating at 4 separate promoter regions and 4 antisense promoters, one of which (asPro1) has a TSS only 119 bp upstream of the Pro1 sense transcript initiation site, indicating the presence of a putative switch element (Figure 5A). The other two GATA3 antisense transcripts are not in close proximity to other promoter elements, however they may function to silence Pro2 and Pro 3, since the last exon of the asPro3 and asPro4 transcripts overlap the Pro2 promoter, and the asPro2 second exon overlaps the Pro3 promoter.

**Figure 5.**
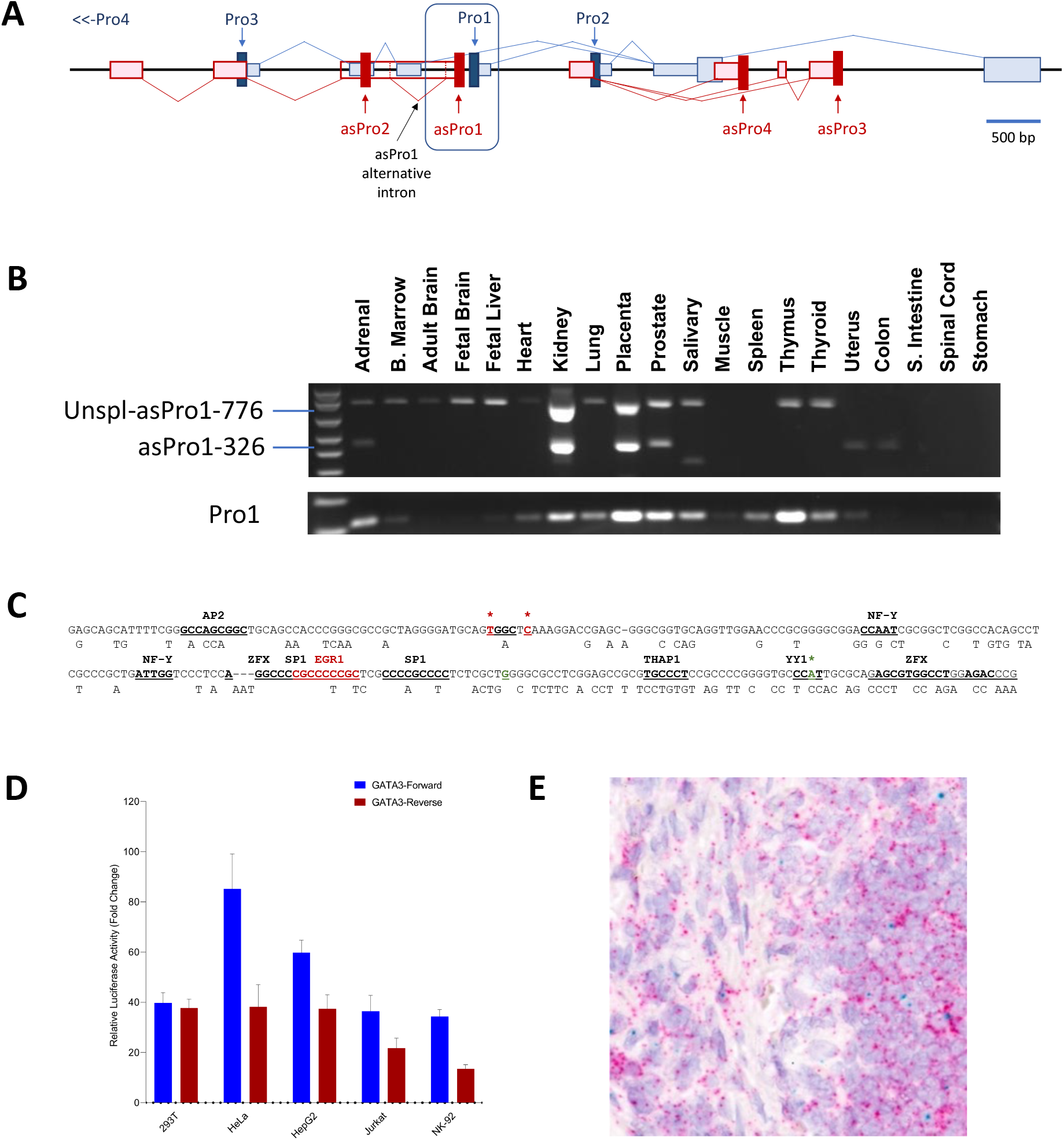
The GATA3 gene has multiple sense/antisense promoters and a bidirectional element. (A) Schematic representation of the 5’ region of the *GATA3* gene is shown. Open blue rectangles represent coding strand exons and larger blue rectangles indicate the open reading frame. Red rectangles depict antisense exons. Exon splicing patterns are indicated by lines joining the exons. The location of an alternatively spliced intron in the asPro1 transcript is indicated by the black arrow. Filled rectangles indicate promoter regions, with antisense promoters (asPro) labeled in red, and sense promoters (Pro) labeled in blue. An additional GATA3 sense promoter is located 5 kb upstream from the region shown. The boxed region indicates the location of a closely spaced pair of divergent promoters, Pro1/asPro1. (B) A cDNA panel of 20 human tissues was subject to PCR with primers specific for either GATA3 antisense-Pro1 (asPro1; upper panel) or coding strand transcripts originating from Pro1 (lower panel). (*C*) The sequence of the human *GATA3* gene in the region spanning divergent Pro1 sense and asPro1 TSS is shown. The coding strand TSS is indicated by the green asterisk, and the pair of antisense TSS are marked by red asterisks. Predicted TF-binding sites are indicated by bold underlined type. Only differing nucleotides present in the mouse *GATA3* gene are shown below the human sequence. (*D*) The relative luciferase activity of a 204 bp *GATA3* bidirectional promoter fragment in five cell lines is shown. Fold change in light units relative to empty pGL3 vector is shown on the y-axis. Values represent the mean and error bars indicate the SEM of at least three independent experiments. Promoter activity from either GATA3-Forward (Pro1; coding strand) or GATA3-Reverse (asPro1) orientations of the bidirectional promoter fragment are shown for each cell line. (*E*) In situ RNA hybridization with RNAscope probes was performed on human thymus tissue. The red spots indicate binding of a probe that recognizes a 118 bp region in the first exon of the *GATA3* Pro1-antisense transcript. The blue spots correspond to a probe that binds to a 126 bp region contained within the first exon of the Pro1-derived GATA3 mRNA.

RT-PCR with primers specific for either sense or antisense Pro1 transcripts demonstrated the presence of sense transcript in approximately half of the human tissues examined (Figure 5B). Two antisense PCR products were detected due to alternative splicing of an intron (Figure 5A). Sense transcripts were dominant in spleen and thymus, whereas other tissues had an equal or greater level of antisense, such as kidney, placenta, and prostate, indicating switch-like behavior of the asPro1/Pro1 element.

An examination of the Pro1/asPro1 region revealed the presence of four TF-binding sites that were shared with the RORγT bidirectional promoter (AP2, SP1, NF-Y, YY1). In addition, the *GATA3* region contains a pair of predicted ZFX-binding sites, a transcription factor associated with peripheral T cell self-renewal and proliferation (Smith-Raska et al., 2018). However, unlike the *RORγT* bidirectional promoter, there was much lower homology with the mouse *GATA3* gene, and most TF-binding sites are not conserved (Figure 5C).

A 204 bp fragment containing the *GATA3* bidirectional promoter region was cloned into the pGL3 reporter vector in both the sense and antisense orientations. Figure 5D shows the relative promoter activity of the *GATA3* bidirectional promoter in several cell lines. The forward promoter activity was dominant in four of the five cell lines tested, with more balanced activity observed in 293T cells. However, In situ RNA detection of sense and antisense in human thymus tissue demonstrated primarily antisense activity from this promoter region (Figure 5E), which suggests that the 204 bp fragment chosen for luciferase assays lacked key elements supporting antisense transcription. Although few in number, the cells expressing *GATA3* Pro1 transcripts did not express antisense transcripts, consistent with this element behaving as a binary switch.

### The asPro3/asPro4 promoters may act to silence GATA3

The *GATA3* Pro2 promoter appears to represent the major promoter responsible for generating *GATA3* coding transcripts in many tissues (USC Genome Browser; EPD; Figure 6A). The asPro3/asPro4 promoters are located in the second *GATA3* intron and produce a spliced transcript containing 2 or 3 exons. The final antisense exon overlaps with the *GATA3* Pro2 promoter region, reminiscent of an antisense promoter found in the second intron of the human *KIR* genes (Wright et al., 2013). The third exon of the *KIR* antisense transcript overlaps with the proximal *KIR* promoter region, and it was shown to silence KIR expression through a piRNA generated from the promoter region (Cichocki et al., 2010). RT-PCR of human tissue RNAs with primers specific for either *GATA3* Pro2 sense or antisense Pro3 transcripts demonstrated the presence of Pro2 sense transcript in more than half of the tissues examined, and Pro3 antisense was observed in six tissues (Figure 6A). However, unlike the *RORχT* or *GATA3* asPro1 antisense transcripts, which could be detected in the absence of sense transcript in some tissues, the *GATA3* asPro3 transcripts were only detected in tissues with strong expression of Pro2 sense transcripts, suggesting shared regulatory elements.

**Figure 6.**
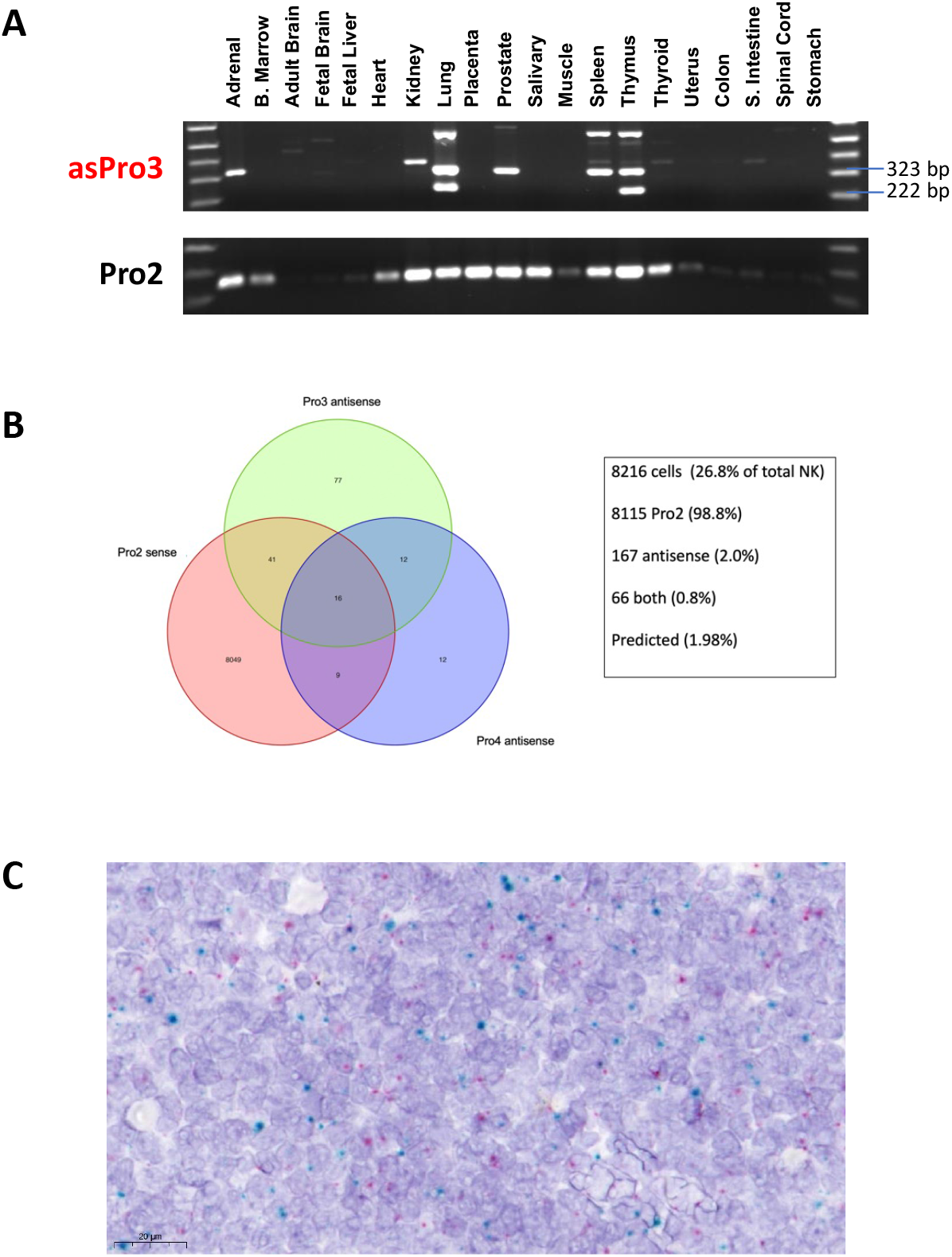
Opposing Antisense-Pro3 and Pro2-sense transcripts are mutually exclusive. (*A*) A cDNA panel of 20 human tissues was subject to PCR with primers specific for either GATA3 antisense-Pro3 (asPro3; upper panel) or coding strand transcripts originating from Pro2 (lower panel). Multiple antisense products are observed due to differential splicing. (*B*) A Venn diagram summarizing the expression of GATA3 sense and antisense transcripts in individual cells is shown. The two major antisense splice forms (Pro3 antisense, Pro4 antisense) are shown together with GATA3 Pro2 sense transcripts. (*C*) In situ RNA hybridization with GATA3-Pro2 and asPro3-specific probes was performed on human thymus tissue. The red spots indicate binding of a probe that recognizes a 130 bp region in the first exon of the Pro3 *GATA3* antisense transcript. The blue spots correspond to a probe that binds to a 201 bp region contained within the first exon and 12 bp at the start of the second exon of the Pro2-derived GATA3 mRNA.

The TSS of the opposing Pro2 and asPro3 promoters are separated by approximately 1.5 kb, unlike the closely spaced divergent transcripts of the bidirectional promoter regions. Therefore, it was of interest to determine if they could be co-expressed in the same cell.

Bioinformatic analysis of human peripheral blood NK single cell RNA seq revealed that ~27% of NK cells express GATA3-coding transcripts that initiate from Pro2. Antisense transcripts from either asPro3 or asPro4 were only detected in 0.5% of the NK cells. The co-expression of GATA3 sense and antisense transcripts is depicted in Figure 6B. Of the cells expressing GATA3 transcripts, 99% expressed sense, and only 2% expressed antisense. Cells expressing both sense and antisense accounted for only 0.8%, consistent with independent expression of sense and antisense transcripts.

The potential independent/mutually-exclusive expression of sense and antisense was confirmed by in situ hybridization with Pro2 and asPro3 specific probes performed on human thymus tissue, where strong expression of both sense and antisense were detected (Figure 6A). Similar levels of Pro2 sense and asPro3 transcripts were detected, and they were rarely co-expressed, indicating mutually exclusive promoter activity (Figure 6C). This result provides further proof that that opposing promoters can also function as a switch element.

## Discussion

It is well accepted that it is the timing and expression of transcription factors that determine the pathways of cell differentiation during development. The actual mechanisms that program development may be analogous to the Boolean logic gates that have been used to program computers. The paradigm of logic gates controlled by multiple inputs has been applied to the complex interactions of TFs for some time (Struhl, 1999), however, the ability of individual TF promoters to behave as binary switches adds a new dimension to the concept of biological logic gates. One can consider these binary promoter switches as logic gates, and the complexity of each gate is determined by the number of TF binding sites that modulate the choice of either the “on” or “off” state. Existing algorithms to analyze logic gates (Yan et al., 2019) are generally compatible with the bidirectional and opposing promoter types of switches described in the *RORC, GATA3*, and *AHR* genes in this study. Transcription factors that stimulate the antisense promoter would be interpreted as negative signals inhibiting gene expression, and would fit into the model. However, if one considers the possibility that the non-coding RNA may in fact have an effect on gene expression, then there are three possible states: sense, antisense, and “off”. In addition, the potential probabilistic behavior of these elements together with the observation that sense and antisense transcription can co-exist in the same cell, further complicates modeling of the system. Finally, it is important to consider that in many cases, there is not a single promoter governing the expression of a single TF gene, such as the eight separate promoters observed in the GATA3 gene, which suggests the presence of cell typespecific switches.

The question of why the two types of binary elements exist within a given TF gene, may be related to the amount of time a given gene will spend in the “on” or “off” states. Bidirectional promoters would have the capacity to quickly switch between states, since a common control region is maintained in an active state. This may also provide an explanation for the observation of promoters in a “poised” state, where the chromatin has the attributes of an active promoter, but no coding transcripts are detected. The opposing promoter type of switch may produce more stable “on” or “off” states due to the epigenetic silencing of the downstream opposing promoter. The probabilistic behavior of bidirectional promoters together with the well-established phenomenon of transcriptional bursting (Rodriguez et al., 2020), could result in relatively long periods where TF mRNAs are absent, creating significant variegated expression of key lineage-determining transcription factors. This may be an important element contributing to the multilineage priming observed in progenitor cell populations (Zhou et al., 2019; Olsson et al., 2016; Mercer et al., 2011). If one considers the two TFs involved in a binary fate choice between the granulocyte (*Gfi1*) and monocyte (*Irf8*) lineages, where there is a metastable mixed-lineage transcriptional state involving the low-level expression of both *Irf8* and *Gfi1* (Olsson et al., 2016), the relative levels of these competing TFs may be modulated by binary switch elements. Examination of the murine *Gfi1* gene in the USC genome browser and EPD, reveals the presence of both bidirectional and opposing antisense promoters. The murine *Irf8* gene does not have any documented antisense transcripts: however, CAGE data indicates an antisense TSS 400 bp upstream from the *Irf8* TSS. Similarly, the human *IRF8* gene shows an antisense TSS 150 bp upstream from the *IRF8* TSS, and a predicted antisense transcript.

To further explore the possible enrichment of divergent promoters in TF genes, the complete set of predicted human sequence-specific transcription factors (Lambert et al., 2018) was examined for the presence of antisense promoters in close proximity to the TSS of the TF (Figure 7A). Of the 1640 likely sequence-specific TFs examined, 33% had divergent antisense TSS within 1 kb, indicating an enrichment relative to the approximately 12% of human genes that have antisense TSS within 1kb (Trinklein et al., 2004). However, in contrast to the analysis of all human genes, which demonstrated a peak of divergent TSS separation around 300 bp, associated with coordinated gene regulation, the TFs produced a peak spanning 70-200 bp (Figure 7A), indicating possible promoter competition generating switch elements. Antisense transcripts opposing the TF promoter were also frequently found, comprising 30% of the TFs. The majority of the opposing transcript TSS were found within 1.5 kb of the gene TSS (Figure 7B), consistent with their localization in the first intron of the gene. Remarkably, 18% of TFs possessed both divergent and convergent promoter pairs, as observed for the three lineage-determining TFs examined in the current study. This suggests that a common promoter structure exists for lineage-determining TFs (Figure 7C), where a bidirectional element creates variegated expression of the TF in progenitor cells, and an opposing transcript silences TF expression if it is not a lineage-defining TF for the cell fate chosen. Although it is highly likely that most opposing promoters will silence gene expression, examination of expression patterns by single-cell RNAseq or *in situ* hybridization in relevant cell types is required to determine if there is competition or coordinated regulation of divergent transcripts with TSS within 300 bp.

**Figure 7.**
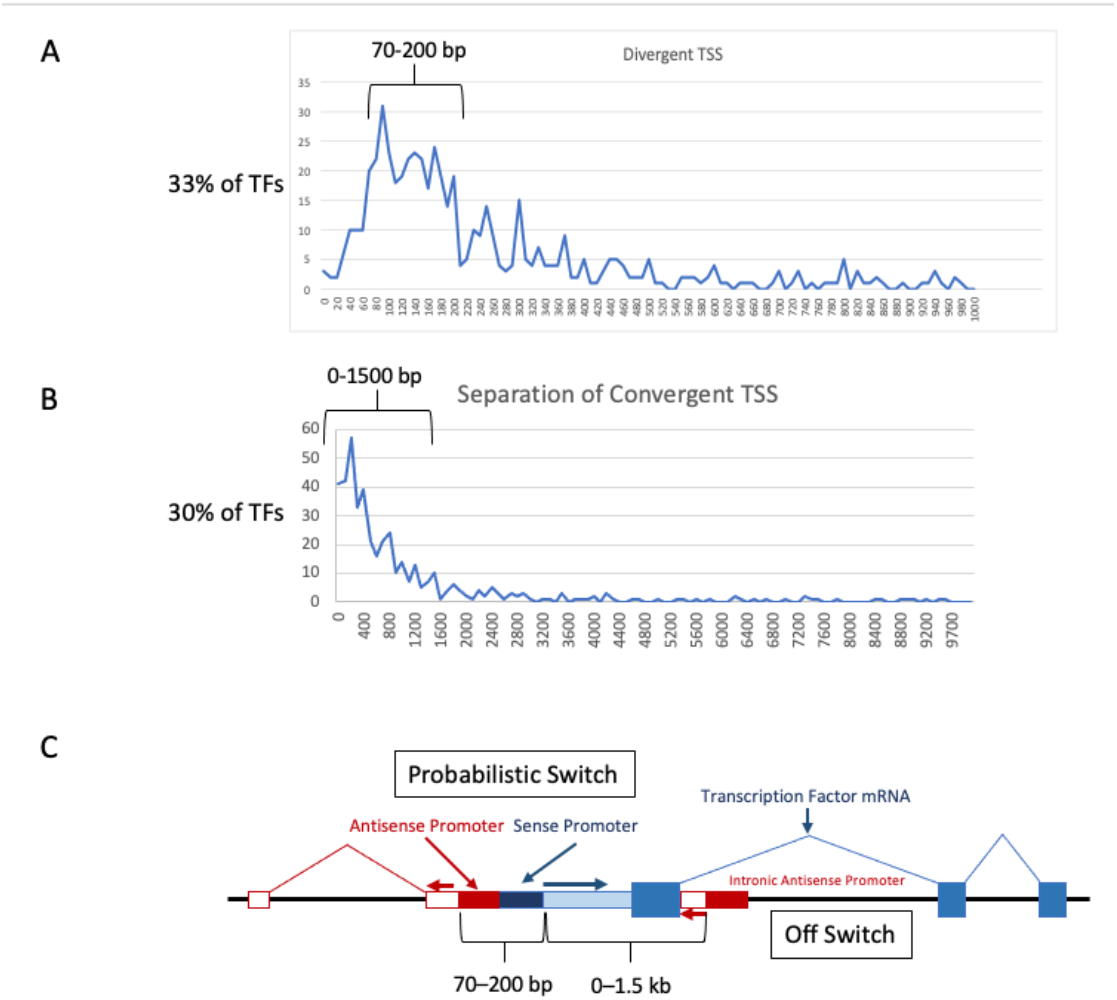
Frequent presence of both divergent and convergent promoters in human transcription factors. A summary of separation distances of antisense TSS from the major TSS of human TFs. (A) shows the profile of TFs that have divergent antisense transcripts initiating within 1 kb of the coding stand TSS, which comprise 33% of all TFs. The enriched 70–200 bp range of TSS separation is indicated. (B) shows the profile of TFs that have a convergent antisense transcript within 10 kb of the coding strand TSS, which comprise 30% of all TFs. The enriched 0-1500 bp region is indicated. (C) shows the general structure of TFs controlled by both bidirectional (Probabilistic Switch) promoter pairs and opposing antisense transcripts (Off Switch), which comprise 18% of all TFS.

In addition to the switch behavior generated by competing promoters, there are likely functional consequences related to the expression of the non-coding RNAs generated by these switch elements. The *GATA3-AS1* transcript expressed from the asPro1 or asPro2 regions (Figure 5A) was shown to be necessary for the efficient expression of the *GATA3, IL5*, and *IL13* genes (Gibbons et al., 2018). The proposed mechanism of action involved recruitment of the MLL methyltransferase responsible for H3K4 methylation, and R-loop formation to tether MLL to the locus. Two recent studies of *SPI1*(*PU.1*) regulation in AML revealed functional roles for a non-coding RNA transcribed from an antisense promoter (asRNA) in the 3rd exon of the gene, and a lnc-RNA termed LOUP that is transcribed from a promoter/enhancer element 17 kb upstream from the *SPI1* promoter (Trinh et al, 2021; van der Kouwe et al., 2021). The asRNA was shown to reduce *SPI1* mRNA, and the LOUP lnc-RNA facilitated interaction of the upstream enhancer with the *SPI1* promoter and stimulated expression. RUNX1 regulates the production of these RNAs, and the RUNX1-ETO fusion protein favored asRNA expression, thus blocking myeloid differentiation. The authors proposed that the competitive interaction of the upstream enhancer with either the proximal or antisense promoter creates a binary on/off switch for either myeloid (SPI1 on) or T cell development (SPI1 off).

In conclusion, we have uncovered a novel mechanism contributing to the programming of cell fate through differential transcription factor expression: the presence of both dynamic and static promoter switching mechanisms that can generate variegated expression patterns of lineage-determining transcription factors in progenitors, creating metastable mixed-lineage states.

## Materials and Methods

### Cell Culture

HeLa and HEK293T cells were cultured in Dulbecco’s modified Eagle’s medium and HepG2 cells were cultured in Eagle’s minimum essential medium. The Jurkat cell line was maintained in RPMI1640 Medium. Each culture medium contained 10% fetal bovine serum (FBS), 100 U/ml each of penicillin and streptomycin (P/S), sodium pyruvate, and L-glutamine. The NK-92 cell line was cultured in α-minimum essential media (ThermoFisher, Waltham MA, USA) containing 0.1 mM beta-mercaptoethanol, 0.02mM Folic Acid, 20% FBS, 100 U/ml P/S, and 200 U/ml of recombinant human IL-2 (Roche, Nutley NJ, USA). All cell lines were grown in cell culture incubators at 37°C in the presence of 5% CO2.

### NK cell isolation

Healthy volunteers were recruited through the NCI-Frederick Research Donor Program (http://ncifrederick.cancer.gov/programs/science/rdp/default.aspx). NK cells were separated from the peripheral blood of healthy donors by Histopaque (Sigma-Aldrich, St Louis, MO, USA) gradient centrifugation using the RosetteSep Human NK Cell Enrichment Cocktail (STEMCELL Technologies, Vancouver, BC, Canada).

### Sorting of CD56^+^ NK Cells

Human NK cells purified from peripheral blood were washed two times in sorting buffer containing 1X PBS pH 7.4, 1% FBS, 25 mM HEPES and 1mM EDTA and pelleted by spinning at 200 RCF for 10 minutes. The samples were resuspended in 200 μl and stained on ice 30 minutes with APC conjugated anti-CD56 antibody (Beckman Coulter) and subsequently washed. CD56^+^ bright and dim cells were sorted into NK media (RPMI 10 % FBS, 100 U/ml penicillin and 100 μg/ml streptomycin, 0.29 mg/ml L-glutamine, 1mM sodium pyruvate, 10 mM HEPES, 0.1 mM B-mercaptoethanol and 500 U/ml human recombinant IL-2 and allowed to recover 24 hours prior to further analysis.

### Single Cell RNA sequencing

Single NK cell suspensions were purified from the blood of four donors. NK cells were washed with PBS + 0.04% BSA at room temperature, resuspended in 500-1000 ul of the same buffer and counted. The cell viability of the samples was 93%-96%. Approximately, 10,000 cells were loaded on the 10X Genomics Chromium platform and libraries were constructed with a Chromium Next GEM Single Cell 5’ reagent kit v2 dual index according to the manufacturer instructions. The libraries were sequenced on NovaSeq S1 100 cycles asymmetric paired-end run with read length of 10 bp for the sample index reads and 150 bp for the Read 1 and Read 2 to achieve a sequencing saturation about 70%.

### Bioinformatics Methods

Quantification of the single cell transcripts was performed with Alevin tool in Salmon software (version 1.5.2; Srivastava et al., 2019). Refseq GRCh38.p13 (accession GCF_000001405.39) assembly transcripts with antisense AHR and GATA3 transcripts included was used as the reference. Seurat (V4) R package was applied for the downstream single cell transcript analysis (Hao et al., 2021). Cell Ranger v6.0.2 analyzed cells were QC filtered (nCount_RNA < 40000 & nFeature_RNA > 400 & nFeature_RNA < 5000 & percent.mt < 10) to remove dead and low-quality cells. Filtered cells were extracted from the Salmon/Alevin quantification for normalization, clustering, visualization as well as differential expression (DE) analysis.

Data normalization performed with Seruat v4 NormalizeData() function. Clustering performed with 35 principal components explaining most variance selected based on their rankings displayed by Seurat’s Elbowplot() function. FindNeighbors() function was run to generate a nearest neighbor graph with the dims=1:35 as dimensions of reductions with the default settings. Clusters of cells were identified by a shared nearest neighbor (SNN) modularity optimization based algorithm by utilizing the FindClusters() function at 0.3 resolution. Clusters are visualized with Uniform Manifold Approximation and Projection (UMAP) dimensional reduction technique with 35 PCs.

Cells expressing AHR sense only or antisense only were selected and differential expression analysis was performed for each group compared to all other cells with Seurat’s FindMarkers() function with default settings. Identified markers were filtered for Bonferroni corrected adjusted p-value <0.05. Differentially expressed transcription factors were identified from the human transcription factors database downloaded from http://humantfs.ccbr.utoronto.ca/index.php.

### Human tissue cDNA panels and RT-PCR

A panel of 20 normal human tissues was purchased from OriGene Technologies (Rockville, MD, USA). RNA quality and quantity were determined using Agilent RNA 6000 Nano Chip (Agilent Technologies Inc., CA, USA). cDNA synthesis was carried out using Random Hexamer primer, Taqman Reverse Transcription Reagents kit (Applied Biosystems, Foster City, CA, USA) according to the manufacturer’s instructions. The RT-PCR reactions were performed in 25μl final volume containing 20 ng of cDNA, 1 x ZymoTaq PreMix (Zymo Research, Irvine CA, USA) and 10μM of each primer, respectively. The PCR conditions were as follows: 95°C for 10 min, 38 cycles of 95°C for 15 sec, 58°C for 45 sec, and 72°C for 20 sec. All primers used in this study were listed in Table S1.

### Generation of luciferase reporter plasmids

Promoter fragments from AHR, GATA-3 and RORC were generated by PCR with ZymoTaq PCR Master Mix (Zymo Research) using the primers listed in Table 1. Inserts were ligated using GenBuilder™ Cloning Kit (Genscript, Piscataway NJ, USA) into pGL3-Basic (Promega, Madison WI, USA) vector linearized with XhoI/HindIII (NEB, Ipswich MA, USA) to generate constructs in both forward and reverse orientations. Each construct was verified by Sanger sequencing. Plasmid DNA for each verified construct was isolated using ZymoPURE II Plasmid Midiprep kit (Zymo Research).

### Cell transfection and Luciferase Reporter Assay

HeLa, HEK293T and HepG2 cells were plated at 2.0×10^5^ cells per well in a 24-well plate the day before transfection and incubated overnight at 37°C in 5% CO2. For each well, 400 ng of pGL3 reporter construct plus 100 ng of *Renilla* luciferase pRL-SV40 control DNA was diluted in 100 μl of buffer. 1 μl of jetOPTIMUS (Polyplus, New York NY, USA) transfection reagent was added and incubated at room temperature for 10 min. The DNA mixture containing was then added to each well and incubated at 37°C in 5% CO2 for 48 hours before analysis.

For suspension cell line transfection, 5×10^5^ Jurkat cells were seeded per well in a 24-well plate and 10.0×10^5^ NK-92 cells were seeded per well in a 6-well plate the day before transfection. For each well, 4 μg of reporter construct DNA, 400 ng of *Renilla* DNA, 5 μl of jetOPTIMUS transfection reagent were diluted in 200 μl buffer and incubated at room temperature for 10 minutes before addition to the well.

Luciferase activity was assayed using the Dual-Luciferase Reporter Assay System (Promega) according to the manufacturer’s instructions. Briefly, the culture medium was removed 48 hours post-transfection and cells were washed with 0.5 mL of phosphate buffered saline (PBS, pH 7.4). Cells were then lysed with passive lysis buffer. The suspensions were centrifuged at 14000 rpm for 1 min. A total of 20 μL of cell lysate supernatant was mixed with 100 μL of luciferase substrate, and the light units were measured on a luminometer. Measurement of the firefly luciferase activity of the promoter constructs was normalized relative to the activity of the *Renilla* luciferase produced by the pRLSV40 control vector and each construct was tested in duplicate in at least three independent experiments.

### Single-molecule RNA in situ hybridization

RNA in situ hybridization experiments were performed using the RNAscope™ technology, which has been previously described (Wang et al., 2012). Paired double-Z oligonucleotide probes were designed against each target RNA using custom software. The probes used are shown in Table 2 below.

**Table 2.** RNAscope probes.

| Catalog Number | Probe Name | Probe Symbol | Probe Genbank Accession Number | # Of ZZ Probe Pairs | Target Region |
| --- | --- | --- | --- | --- | --- |
| 1061421-C1 | *Homo sapiens* aryl hydrocarbon receptor (AHR) mRNA | BA-Hs-AHR-3zz-st | NM_001621.5 | 3 | 2-277 nt |
| 1085361-C2 | *Homo sapiens* aryl hydrocarbon receptor (AHR) cDNA clone mRNA sequence | BA-Hs-AHR-O2-3zz-st | BU853219.1 | 3 | 379-527 nt |
| 1084911-C1 | *Homo sapiens* GATA binding protein 3 (GATA3) transcript variant X1 mRNA | BA-Hs-GATA3-O2-3zz-st | XM_005252442.2 | 3 | 32-157 nt |
| 1104171-C2 | *Homo sapiens* GATA3 antisense RNA 1 (GATA3-AS1) transcript variant 3 long non-coding RNA | BA-HS-GATA3-AS1-3zz-st-C2 | NR_104336.1 | 3 | 17-134 nt |
| 1085341-C1 | *Homo sapiens* GATA binding protein 3 (GATA3) transcript variant 2 mRNA | BA-Hs-GATA3-O4-3zz-st | NM_002051.3 | 3 | 4-216 nt |
| 1084931-C2 | PREDICTED: *Homo sapiens* uncharacterized LOC107984204 ncRNA | BA-Hs-GATA3-AS1-O1-3zz-st-C2 | XR_001747358.1 | 3 | 405-534 nt |
| 860431-C1 | *Homo sapiens* RAR related orphan receptor C (RORC) transcript variant 2 mRNA | BA-Hs-RORC-tv2-2zz-st | NM_001001523.2 | 2 | 43-146 nt |
| 860421-C2 | *Homo sapiens* cDNA clone THYMU2002838 5' mRNA sequence | BA-Hs-RORC-intron2-3zz-st-sense | DB103232.1 | 3 | 2-140 nt |

The BaseScope Duplex Reagent Kit (cat. no. 323800) (Advanced Cell Diagnostics, Newark, CA) was used according to the manufacturer’s instructions. FFPE tissue sections or cytospin slide samples were prepared according to manufacturer’s recommendations. Target retrieval and protease pretreatment conditions were optimized for each sample type. Each sample was quality controlled for RNA integrity with probes specific to the housekeeping genes *PPIB* and *POLR2A*. Negative control background staining was evaluated using a probe specific to the bacterial *dapB* gene. Brightfield images were acquired using a 3D Histech Panoramic Scan Digital Slide Scanner microscope using a 40x objective.

## Acknowledgements

This project has been funded in whole or in part with Federal funds from the Frederick National Laboratory for Cancer Research, National Institutes of Health, under contract HHSN261200800001E. This research was supported in part by the Intramural Research Program of NIH, Frederick National Lab, Center for Cancer Research. We would also like to thank the CCR Sequencing Facility at Frederick National Laboratory for Cancer Research for performing the single cell capture and the sequencing.

